# Certain spatial prediction decreases the rhythm of attentional sampling

**DOI:** 10.1101/2023.11.20.567760

**Authors:** Yih-Ning Huang, Wei-Kuang Liang, Chi-Hung Juan

**Affiliations:** Institute of Cognitive Neuroscience, National Central University, No.300, Jhongda Rd, Jhongli, 320, Taiwan; Cognitive Intelligence and Precision Healthcare Research Center, National Central University, Taiwan

**Author notes:** Corresponding author email address Chi-Hung Juan.

## Abstract

Recent studies demonstrate that behavioral performance during visual spatial attention fluctuates at theta (4-8 Hz) and alpha (8-16 Hz) frequencies, linked to phase amplitude coupling (PAC) of neural oscillations within the visual and attentional system. Moreover, previous research suggests that attentional sampling rhythms are task-dependent, evidenced by varying behavioral performance at different frequencies. To investigate the role of prior spatial prediction, we employed an adaptive discrimination task with variable cue-target onset asynchronies ranging from 300 ms to 1300 ms in steps of 20 ms, while manipulating spatial prediction via cue validity (100% & 50%), with concurrent Electroencephalography (EEG) recording. We applied adaptive data analytical methods, namely Holo-Hilbert Spectral Analysis (HHSA) and Holo-Hilbert Cross-frequency Phase Clustering (HHCFPC). Our findings indicate that response precision for near-threshold Landolt rings fluctuates at the theta-band (4 Hz) under certain predictions and at alpha & beta bands (15 & 19 Hz) with uncertain predictions. Furthermore, spatial prediction strengthens theta-alpha modulations at parietal-occipital areas, frontal theta phase and parietal-occipital alpha amplitude coupling, and within frontal theta phase/ alpha amplitude coupling. Notably, during the pre-target period, beta-modulated gamma oscillations in parietal-occipital areas predict response precision in spatially uncertain conditions, while frontal theta phase and parietal-occipital alpha amplitude coupling predict response precision in spatially certain conditions. In conclusion, our study not only strengthens the notion that the speed of periodic sampling in perception depends on the task at hand but also highlights the critical role of spatial prediction in attentional sampling rhythms.

**Significance Statement:** This study investigates the temporal dynamics of sustained spatial attention under varying certainty levels, employing behavioral and electrophysiological measures in an adaptive discrimination task. Unveiling the rhythmic nature of sustained attention, our findings showcase substantial effects of spatial certainty on attentional rhythms, witnessing an increased certainty that shifts these rhythms from beta to theta frequencies. Neural oscillations offer insights into the underlying mechanisms, revealing theta-alpha coupling and beta-gamma coupling within the visual system and frontal-parietal network. Significantly, our results challenge conventional notions of attentional rhythms, emphasizing the dynamic complexity of these processes. In a broader context, our study contributes to bridging the gap between task demands and periodic sampling rhythms, offering novel insights into attention allocation during complex tasks.

## 1. Introduction

Recent studies have presented behavioral and physiological evidence suggesting that sustained attention in the face of uncertain spatial and temporal predictions follows a rhythmic pattern (Fiebelkorn and Kastner 2019; VanRullen 2016). Spatial cueing tasks utilizing fine-grained and variable cue-target onset asynchronies (CTOA) have revealed a trend-based variation in behavioral performance, accompanied by rhythmic fluctuations spanning from the alpha-band (8-16 Hz) to the theta-band (4-8 Hz; Kienitz et al. 2022). It’s noteworthy that the speed of phasic alternations appears to be task-dependent (Dugué and VanRullen 2017; Michel et al. 2022). For instance, rhythmic sampling toward two objects (visual fields or features) was reported in the theta band (4 Hz), while rhythmic sampling towards one was observed in the alpha band (8 Hz; Fiebelkorn et al. 2013; Holcombe and Chen 2013; Landau and Fries 2012; Re et al. 2019).

Prior investigations have consistently demonstrated the existence of rhythmic structures in sustained attention and their correlation with functional connectivity across the frontal-parietal network (i.e., the attentional network) and the visual cortex. Specifically, alpha rhythms exhibit a correlation with alpha phase/gamma amplitude coupling, serving as a natural periodic filter across the brain (Busch and VanRullen 2010; Mathewson et al. 2011; Osipova et al. 2008). Likewise, theta rhythms are linked to theta-phase/alpha-gamma amplitude coupling, potentially arising from top-down communication originating in higher-order areas (Fiebelkorn et al. 2018; Helfrich et al. 2018; Landau et al. 2015).

On the contrary, various studies have delved into the certainty of spatial prediction, often manipulated through cue validity (e.g., Chiau et al. 2011; Liu et al. 2010; Posner 1980). Numerous investigations have demonstrated a correlation between the certainty of prior expectations regarding target location and behavioral responses, such as accuracy and reaction time (Liu et al. 2011; Posner 1980; Tseng et al. 2013; Vossel et al. 2014; Vossel et al. 2006), as well as neural correlates during the anticipatory period, including alpha power lateralization (Haegens et al. 2011; Voytek et al. 2017). However, recent studies have presented mixed findings concerning sampling rhythms. While earlier research suggested that spatial weighting might be independent of attentional sampling rhythms (Fiebelkorn and Kastner 2019), recent investigations comparing three different cue validities (i.e., 100%, 80%, or 33%) in a modified Egly-Driver task did not yield substantial behavioral evidence for rhythmic sampling behavior (van Der Werf et al. 2022). Hence, the extent to which rhythmic attention is confined to spatial uncertainty remains unclear.

In the context of brain wave analysis, the importance of precise phase extraction from nonlinear and nonstationary brain signals has been underscored through the application of adaptive analytical methods (Aru et al. 2015; Cole and Voytek 2017; Huang et al. 2016; Hyafil 2015). Studies employing nonlinear decomposition and Hilbert transform, particularly the Holo- Hilbert Spectral Analysis (HHSA), have demonstrated enhanced accuracy in analyzing phase- amplitude modulated brain signals entrained from phase-amplitude coupled external visual stimuli (Juan et al. 2021; Nguyen et al. 2019). Additionally, adaptive neural correlates optimizing behavioral performance in attentional tasks, such as inter-site phase coherence (ISPC) and inter-trial phase coherence (ITPC), can be quantified through precise phase extraction (Chang et al. 2023; Liang et al. 2021).

This study aimed to investigate the impact of prior spatial expectations on the sampling mechanism. The examination of behavioral measurements and their underlying neural mechanisms was conducted through an adaptive Posner-like discrimination task, accompanied by EEG recording and adaptive data analytical methods. Our hypothesis posited that uncertain spatial predictive cues (i.e., 50% cue validity) would foster more robust communication between higher associative and visual areas, resulting in theta-band periodic alternations in behavioral performance and theta-phase/alpha amplitude coupling within the attentional system, as observed in previous studies (Helfrich et al. 2018; Plöchl et al. 2022). Conversely, it was anticipated that certain spatial predictions would not induce alternations between objects. However, maintaining a robust spatial-temporal modulation, such as lateralized alpha power, was expected to lead to a more significant improvement in behavioral performance.

## 2. Materials and Methods

### 1.1 Participants

Seventeen participants were recruited from National Central University (10 females and 7 males, mean age = 21.9 years, SD = 1.6 years). All participants had normal or corrected-to-normal visual acuity. One outliner was excluded from the analysis due to poor accuracy in one condition (< 5%). Every participant provided written informed consent prior to taking part in the study. The study protocols received approval from the local IRB committee, specifically the National Taiwan University Research Ethics Committee, under the reference number 201905EM080.

### 1.2 Apparatus and Data Acquisition

The experiment was conducted in a dark and soundproof room, where participants were instructed to sit at a distance of 60 cm from a 54-cm LCD monitor with a vertical refresh rate of 200 Hz. The task was administered using Psychophysics Toolbox Version 3 (PTB-3; Brainard and Vision 1997; Pelli and Vision 1997) in MATLAB (R2019a). Participants placed their heads on the chin-rest and fixated their eyes on the central white cross (138.1 cd/m^2^) on the screen during the whole experiment. EEG signals were concurrently recorded during the experiment through high-density 128-channel sponge-based HydroCel Geodesic Sensor Nets and Net Station software (Electro Geodesics Inc. [EGI], Eugene, OR, USA) with a sampling rate of 1000 Hz. Electrode impedances were evaluated before each block and kept below 50 kΩ.

### 1.3 Experimental Design

Participants were instructed to perform an adaptive orientation discrimination task that was adapted from Michel et al. (2021). The task consisted of blocks with different cue validity (i.e., 100% & 50%) and variable cue-target onset asynchronies (CTOA) ranging from 300 ms to 1300 ms in steps of 20 ms. To ensure a fair comparison in the data analysis, trial numbers in valid trials were identical in each cue validity condition. The experiment included a total of 1500 trials performed over 3 days. Each day consisted of a 100-trial practice session with 50% cue validity only, followed by a main session with 100% and 50% cue validity in five blocks. The cue validity was informed before every session started, and the order of each condition was counterbalanced between participants.

The diagram of one valid trial is illustrated in **Figure 1A**. Every trial was self-initiated with a 300 ms fixation cross (size: 0.7°, 138.1 cd/m^2^) centered in the screen (background luminance: 80.05 cd/m^2^) followed by two placeholders (square line, size: 2.8°, 138.1 cd/m^2^) presented for a random duration of 1800 ms to 2000 ms across trials. The placeholders in the two visual fields indicated the subsequent target position with a visual angle of 3.5° away from the center.

**Figure 1.**
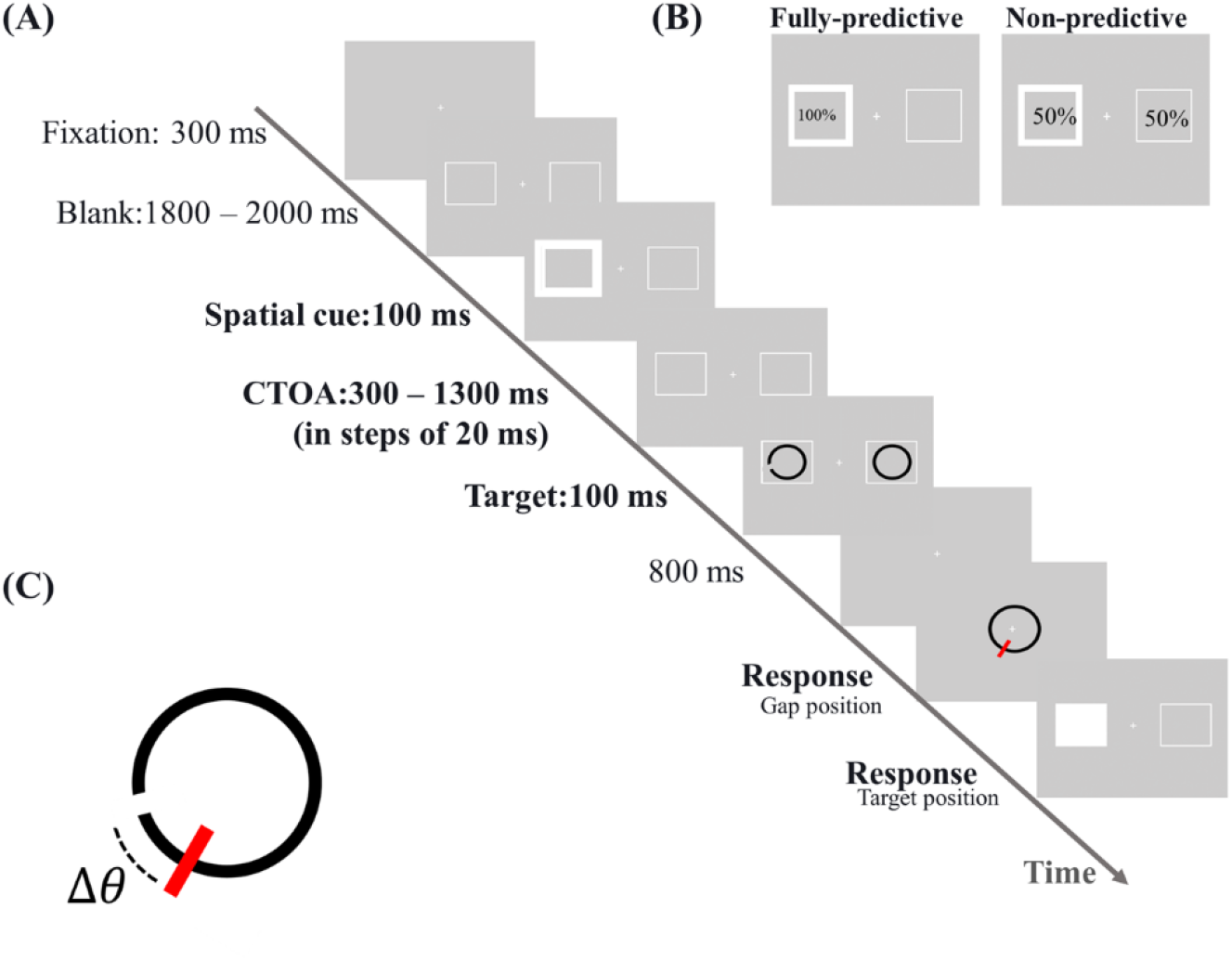
Experimental procedures and conditions (A) An example and the timeline of a valid trial. (B) Certainty of spatial prediction was manipulated by varying cue reliabilities. The cue provided 100% or 50% information about target locations in different blocks. (C) Precision errors were measured by the angular difference between the displayed and reported gap positions, ranging from -180° to 180°.

An exogenous cue (i.e., flash) was then displayed on one side of the placeholder for 100 ms, followed by a variable CTOA pseudo-randomly selected from 300 ms to 1300 ms in steps of 20 ms. In valid trials, the Landolt ring target (size: 1.4°, thickness: 0.175°, gap size: 0.05°) was presented in the placeholder where the cue was previously located, and a closed ring of similar parameters as the Landolt ring was presented in the other placeholder simultaneously for 100 ms. The gap position on the Landolt ring was randomly selected from 0° to 360° across trials.

In invalid trials, the target was presented at the opposite placeholder from the cue.

Following an 800 ms blank interval, there were two response windows: gap and target positions. In the first window, a closed ring (size: 2.8°, thickness: 0.35°, 0.238 cd/m^2^) was presented at the center of the screen. Participants were instructed to move the red bar and click the left mouse button to indicate the position of the gap. The response window would not switch until the answer was received. In the second window, two placeholders identical to the previous stimuli were presented on the screen. Participants were again asked to move their mouse to indicate the target position and click the mouse button when the placeholder was filled to respond. In the practice session, a feedback window was displayed after the response, revealing the response deviation and correctness of the target position on the screen **(Figure 1C)**.

Following previous studies (Landau and Fries 2012; Michel et al. 2022), the target luminance in the study was individually adjusted to maintain performance under 50% cue validity at around 50%. In the practice session, including 100 trials, the luminance of the target and non- target was adjusted using two-up-two-down methods. A trial was deemed correct when the absolute deviation was below 45° and the target position was accurate. The mean luminance of the target was 32.48 cd/m^2,^ with a standard deviation of 3.93 cd/m^2^ between subjects. The mean defined accuracy in the formal test was 60.29% ± 1.79% (mean ± SE) under 50% cue validity and 83.69% ± 2.20% under 100% cue validity condition.

### 1.4 Behavioral Analysis and Statistics

The analysis of the behavioral data was conducted using MATLAB (R2021a). The precision errors were calculated as the smallest angular difference between the displayed and reported gap positions, ranging from -180° to 180°. To assess the effect of cue validity (50% & 100%) and cued visual field (RVF & LVF) on precision errors across all CTOAs, two-way repeated measures ANOVA was performed on averaged precision errors in corresponded conditions.

Another two-way repeated measures ANOVA was conducted to estimate the cueing effect (valid & invalid) under 50% cue validity and the effect of cued visual fields (right & left) on the precision errors.

#### 1.4.1 Behavioral Spectral Analysis

To investigate the dynamic changes in behavior, the deviation was z-score normalized within each participant, day, and conditions, irrespective of the CTOAs, to reduce individual differences. The standardized deviations were pooled across participants within conditions into 50 CTOA bins. For the oscillatory feature in behavioral time series data, spectral analysis was conducted using permutation tests on the power spectrum to reveal the frequency domain of fluctuated deviations. The analytical steps of spectral analysis were similar to those employed in previous studies (Fiebelkorn et al. 2013; Landau and Fries 2012; Michel et al. 2022). In the present study, the linear-detrended grouped time series data was transformed into the frequency domain using Fast Fourier Transform (FFT). The data was applied 50-points discrete Fourier transform (DFT) algorithm using ‘fft’ functions on MATLAB (R2021a; Frigo and Johnson 1998). Permutation tests on the power spectrum were then conducted to test the null hypothesis to ensure no temporal rhythmic structure was present in each condition. To generate random datasets at the individual level, the CTOA labels were shuffled within conditions, resulting in 50000 random time series data sets. These random data sets were then transformed into the frequency domain using the abovementioned steps. Finally, the power values from the original data were compared with those from the 50000 random simulations in each frequency bin ranging from 2 Hz to 20 Hz with a frequency resolution of 1 Hz. The significance level was set at p < 0.05. To control for multiple comparisons of 19 frequency bins, Bonferroni correction was applied, and the p-value threshold was corrected to 0.05/19 = 0.0026.

### 1.5 EEG Analysis and Statistics

#### 1.5.1 EEG Pre-processing

The EEG data were pre-processed with MATLAB (R2020b; R2021a), EEGLAB, and SPM 12 functions (https://www.fil.ion.ucl.ac.uk/spm/) for further analysis. The continuous EEG signals underwent high-pass filtering at 1 Hz, low-pass filtering at 100 Hz, and notch filtering at 59 Hz to 61 Hz. The data was then subjected to channel removal, including facial channels and channels with more than 5 seconds of flat signals, high-frequency noise (> 4 standard deviations to the baseline), and high correlations between nearby channels. Independent component analysis (ICA; Hyvärinen and Oja 2000; Makeig et al. 1996) was applied with second-order blind identification (SOBI; Belouchrani et al. 1997; Belouchrani et al. 1993) to remove components with eye movements, line noise, channel noise, and heart signals. Finally, the EEG signals in each channel were re-referenced to the average of all channels, with the removed brain channels interpolated.

To compare cognitive function without target overlapping in the variable CTOA design, the pre-processed data was segmented into two epochs: cue onset and target onset. The cue onset epochs consisted of the period 1000 ms prior to cue onset to 1300 ms after cue onset, while the target included the time range 1000 ms before target onset to 1300 ms after target onset. The segments underwent artifact rejection to exclude trials with high voltages (> 200μV).

#### 1.5.2 Holo-Hilbert Spectral Analysis (HHSA)

Previous research emphasized the importance of coupling between different frequency bands, including cross-frequency coupling (CFC), phase amplitude coupling (PAC), and amplitude modulations (AM), in cognition (Canolty and Knight 2010). However, linear algorithms have limitations in accurately analyzing nonstationary and nonlinear brain signals, particularly the instantaneous phase (Aru et al. 2015; Cole and Voytek 2017). Therefore, the current study utilized the nonlinear Holo-Hilbert Spectral Analysis (HHSA) method, with the first step of the multiple-layer Complete Ensemble Empirical Mode Decomposition with Adaptive Noise (CEEMDAN) method, to analyze the preprocessed EEG segments to enhance the understanding of neural oscillations (Chang et al. 2023; Huang et al. 2016; Juan et al. 2021; Liang et al. 2021).

The adaptive decomposition algorithm, or CEEMDAN (Torres et al. 2011), is an improved version of the empirical mode decomposition (EMD; Huang et al. 1998) and ensemble empirical mode decomposition (EEMD; Wu and Huang 2009). The EMD decomposes the signal into multiple Intrinsic Mode Functions (IMFs), with each IMF satisfying two principles:

(1) The difference between the number of relative extrema and zero-crossings is less than or equal to one, and (2) the average of upper and lower envelopes is near zero. When the IMF adheres to two principles, the IMF will be subtracted from the residual. IMFs were addressed from high to low frequency, calculated from repeated subtraction of residuals and IMFs. In recent years, to address the issue of mode mixing from the original EMD, the EEMD was expended to add Gaussian white noise of finite variance to the original signal over multiple trials, and subsequently averaged into IMFn from all ensembles to obtain each IMF. This approach helps to reduce the mode mixing problem and improve the accuracy of the decomposition (Huang et al. 1998; Wang et al. 2012). Moreover, to increase the spectral separation from signals decomposed with EEMD, CEEMDAN averages the ensembles of IMF with white noise before subtracting from the residual in each step.

In the current study, the pre-processed EEG data underwent the multiple layer CEEMDAN method (Huang et al. 2016). The first layer decomposed the data into several IMFs (**Figure 2A)** with a standard noise deviation of 0.2 and an ensemble size of 50, given by the equation (1).

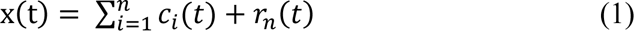

**Figure 2.**
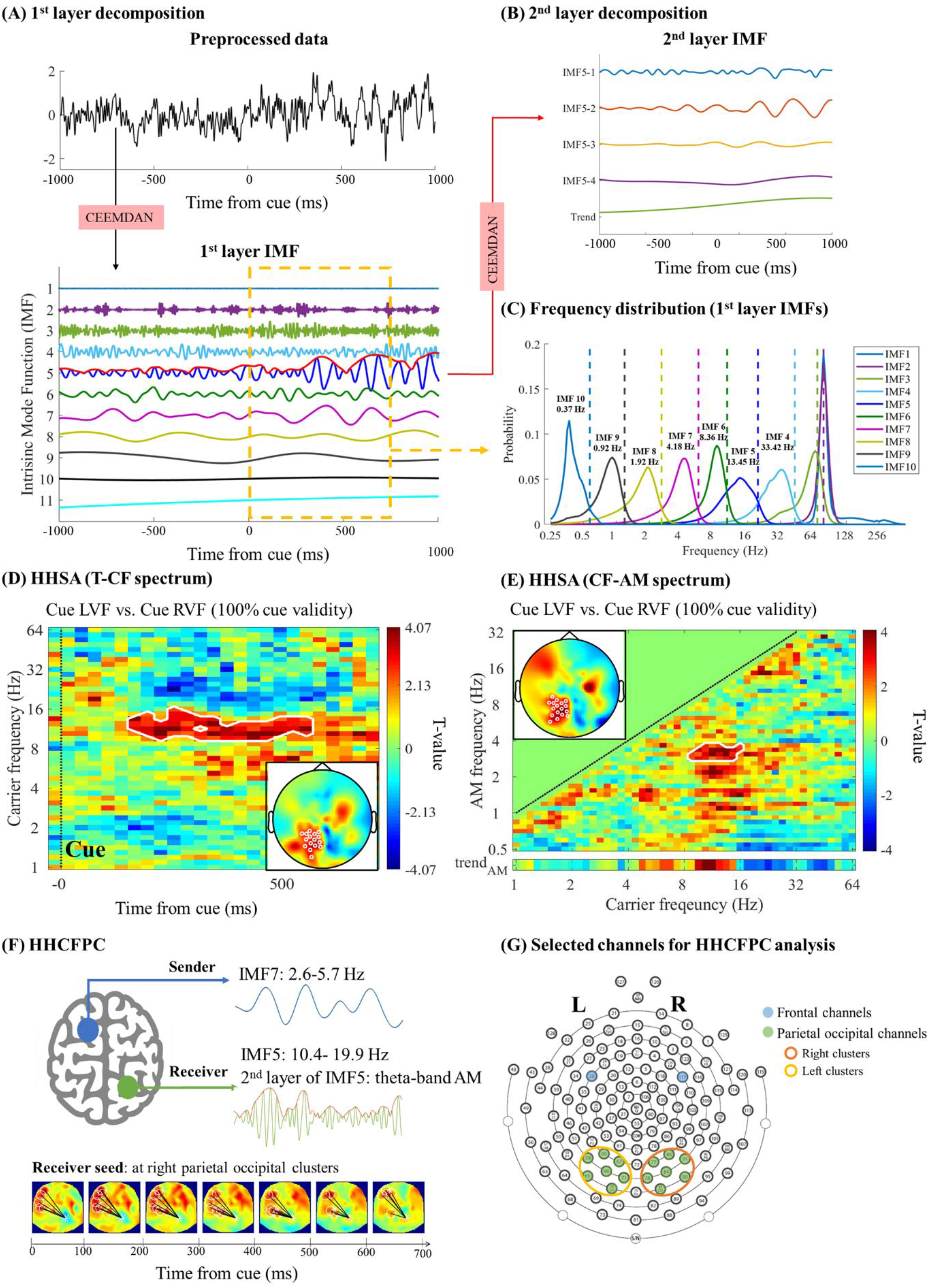
Illustration of multiple layer CEEMDAN method. (A) The pre-processed data underwent decomposition using the complete ensemble empirical mode decomposition with adaptive noise (CEEMDAN) method, resulting in several intrinsic mode functions (IMFs). The amplitude envelope of each IMF was subsequently decomposed into IMFs. (B) illustrates the second layer IMFs of IMF5. (C) depicts the frequency distribution of each first layer IMF within the time range of 0-700 ms following cue onset. (D) Depiction of the 2D Holo-Hilbert time-carrier frequency (T-CF) spectrum, and (E) the carrier frequency-amplitude modulation (CF-AM) spectrum averaged across selected electrodes (time window: 0 to 700 from the onset of the spatial cue). The inserted figure displays the topographic distribution of the statistical results. The white markers indicate significant differences detected by the paired t-CBnPP test (two-tailed, p < 0.05) (F) An HHCFPC example demonstrates theta phase/alpha amplitude coupling between the frontal and parietal channels. The HHCFPC was calculated according to the blue line (IMF5’s second layer IMF phase function) and orange line (IMF7’s phase function). (G) The electrodes selected for the HHCFPC analysis are displayed. (Blue: frontal channels, Green: parietal-occipital channels, Orange: right parietal occipital clusters, Yellow: left parietal occipital clusters)

The preprocessed data was then segmented into different IMFs (*C*_*i*_(*t*) in the above equation), which corresponded to distinct and less overlapping frequency bands and an intrinsic trend without oscillatory features **(Figure 2C)**. Next, the direct quadrature (DQ) method was used to derive the instantaneous phase function for each IMF, and the cubic spline algorithm was employed to obtain the amplitude function (Huang et al. 2009), as represented by the equation

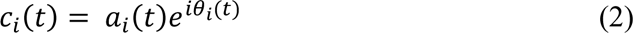

The amplitude mode function *a*_*i*_(*t*) of each first layer IMF was then decomposed through CEEMDAN with a noise standard deviation of 0.2 and an ensemble size of 20, which served as the second layer IMF **(Figure 2B)** in this study. Following the same steps as the first layer, the instantaneous phase and amplitude functions were derived for each second layer IMF using the general zero-crossing methods and cubic spline algorithm, respectively, as represented by the equation (3).

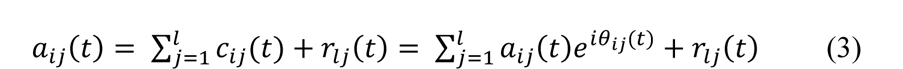

Accordingly, the time information, the first-layer IMFs (i.e., carrier frequency), and the second-layer IMFs (i.e., amplitude modulation frequency) were displayed in a three-dimensional Holo- Hilbert spectrum (HHS) at the same time. However, the statistical comparison between different conditions was conducted on the two-dimensional HHS: Time-Carrier frequency (T- CF) spectrum **(Figure 2D)** and Carrier frequency-Amplitude modulation (CF-AM) spectrum (**Figure 2E)**.

This study aimed to investigate the neural response following the spatial cue and preceding the target. To examine the electrophysiological response to the spatial cue onset, we analyzed the 2D spectrum, consisting of T-CF and CF-AM, across different conditions (i.e, cue validity x cued location). The T-CF spectrum covered the 0-700 ms period after the spatial cue onset. To enhance the normality of spectral power, we rescaled the T-CF spectrum to the logarithm of the ratio of spectral power to the averaged power during the time window from -450 ms to -150 ms before the spatial cue onset. Additionally, we averaged the time dimension of the 3D Holo spectrum across the 0-700 ms period after the spatial cue onset to generate the CF-AM spectrum. Also, we rescaled it to the logarithm of the ratio of spectral power to the averaged power during the time window from -450 ms to -150 ms before the spatial cue onset.

To further investigate the neural response preceding the target onset, we also analyzed the 2D spectrum (T-CF & CF-AM) to distinguish between precise and imprecise responses in valid trials within each cue validity condition (50% & 100%). To avoid the foreperiod effect (FP) in specific conditions, we labeled trials as precise or imprecise based on detrended behavioral oscillations and segmented them into good and bad performance phases. The 2D spectrum covered the period before the target onset (-700 – 0 ms) and was also rescaled to the logarithm of spectral power.

#### 1.5.3 Holo-Hilbert Cross-frequency Phase Clustering (HHCFPC)

This study calculated the phase-amplitude coupling through Holo-Hilbert cross-frequency phase clustering (HHCFPC; **Figure 2F**; Liang et al. 2021). The calculation of HHCFPC was adapted from the conventional analysis of inter-site phase coherence (ISPC). In the current study, the coherence was calculated through equation (5):

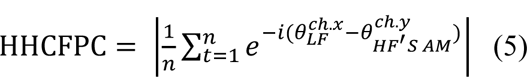

The phase function 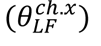 of 1^st^ layer IMF (low frequency, LF) and the phase function 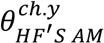 of the 2^nd^ layer IMF (high frequencies’ amplitude modulation envelops, HF’s AM), whose averaged instantaneous frequency matches the LF’s, can be selected at the same (x = y) or between two channels (x ≠ y).

The calculation of HHCFPC in this study was selected based on previous findings, which suggested frontal-parietal networks and occipital areas have significant theta-phase/ alpha- beta-gamma amplitude coupling during sustained attention (e.g., Fiebelkorn et al. 2018; Plöchl et al. 2022). In the analysis of neural correlates to the spatial cue, we first set the receiver seed (i.e., IMF5& IMF6: alpha-band 5.7 -19.9 Hz) at the left (i.e., channels E59, E60, E65, E66, E67, E70, E71) and right (i.e., channels E76, E77, E83, E84, E85, E90, E91) illustrated in the parietal-occipital clusters as shown in **Figure 2G**. Then, the HHCFPC was calculated to the sender (i.e., IMF7: theta-band 2.6-5.7 Hz) in all electrodes in a time window of 300 ms in steps of 30 ms. The following analysis of HHCFPC was based on the significant electrodes in the frontal area (left: E28, right: E117), which were sender (IMF7: 2.6-5.7 Hz) seeds. The time- HHCFPC was rescaled to the logarithm of the ratio of spectral power to the averaged power during the time window from -450 ms to -150 ms before the spatial cue onset. On the other hand, the HHCFPC before the target onset period was calculated from frontal seed sender channels (left: E28, right: E117) to all electrodes. The HHCFPC was analyzed in a time window of 300 ms in steps of 20 ms, from 700 ms before target onset to target onset.

#### 1.5.4 Statistical Analysis

The cluster-based non-parametric permutation (CBnPP) test (Liang et al. 2021; Maris and Oostenveld 2007) was applied to this study for most of the comparisons, which included T-CF and CF-AM spectrum, and HHCFPC. The CBnPP test enhances the validity of statistical significance by incorporating neighboring clusters. The analytical procedure consisted of four parts: performing paired t-tests, clustering significant t-values across multiple dimensions, generating a clustered t-distribution from random data, and comparing the clustered t-value with the random distribution. The statistical analysis began with a paired t-test on measurements between conditions. Next, significant t-values were identified within clusters that were formed based on neighboring channels (within 40 mm), time bins, and frequency bins. These significant t-values were summed to obtain a clustered t-value. To determine the significance of the clustered t-value, the condition label was randomly shuffled, and 5000 random datasets were generated. Each iteration involved analyzing the random data using the same approach as the original data, resulting in distributions of randomized clustered t-values. The original clustered t value was considered significant at a two-tailed alpha level of 0.05 if it exceeded the null hypothesis rejection threshold determined from the randomized clustered t value distribution.

## 3. Results

### 3.1 Behavioral results

#### 3.1.1 Overall Cuing Effects

The mean deviations in the 100% cue validity condition were 27.03° ± 2.43° (mean ± SE) at the right visual field (RVF) and 26.03° ± 2.19° at the left visual field (LVF). For the 50% cue validity condition, the mean deviations were 36.23° ± 3.09° (RVF) and 37.50° ± 2.90° (LVF) in valid trials and 57.64° ± 4.09° (RVF) and 58.09° ± 3.61° (LVF) in invalid trials (**Figure 3**).

**Figure 3.**
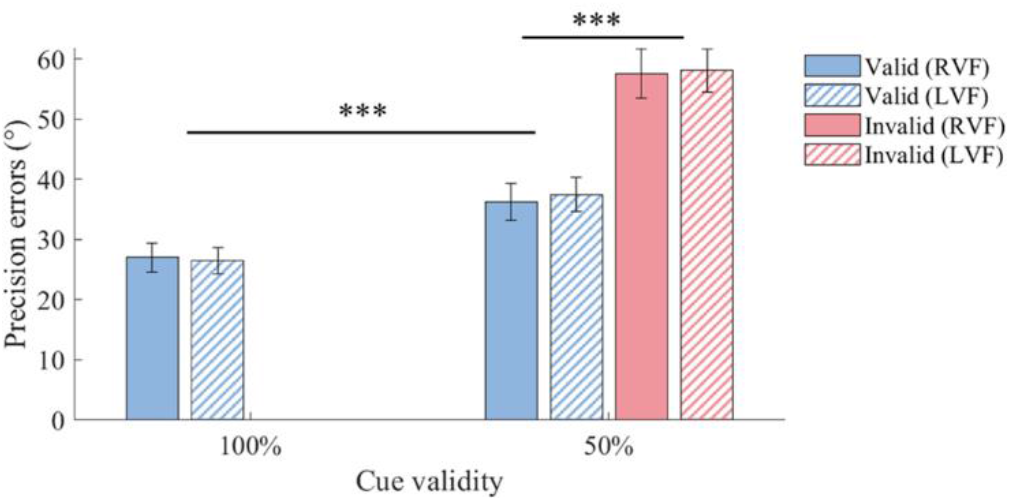
Precision Errors across all CTOAs in Different Conditions. The figure illustrates precision errors for different trial conditions. The blue bar represents valid trials, while the pink bar represents invalid trials. The solid bar corresponds to trials with cues presented in the right visual field, while the slanted bar corresponds to trials with cues presented in the left visual field. The error bar indicates the standard error and *** denotes p< 0.005.

A two-way (VFs x Validity) repeated measures ANOVA was performed on the mean precision errors of valid trials in different cue validity conditions. The results showed that cue validity had a significant main effect (F(1,15) = 33.33, p < 0.001) on deviations in valid trials. meanwhile, no significant effects of cued visual field (F(1,15) = 0.03, p = 0.86) and interaction effects (F(1,15) = 0.32, p = 0.58) were found on the precision errors. Another two-way repeated measures ANOVA was performed on the mean precision errors of valid and invalid trials in the 50% cue validity condition. The results showed that target at valid or invalid locations has main effects on precision errors (F(1,15) = 17.35, p < 0.001). However, there were no effects of visual field (F(1,15) = 0.85, p = 0.37) and interaction effects (F(1,15) = 0.01, p = 0.91) on precision errors.

#### 3.1.2 Temporal Structure of Selective Attention

The temporal pattern of attention in the current task is illustrated in **Figure 4**, which shows the dynamic changes in deviations. After removing the linear trend, fluctuations in standardized precision errors were analyzed. A significant 4 Hz (permutation test, Bonferroni corrected p < 0.05) fluctuation was found in the 100% cue validity condition when the cue was presented in the LVF but not in the RVF **(Figure 5)**. Furthermore, deviations in valid trials fluctuated at 15 Hz (permutation test, corrected p < 0.05) when the 50% informative cue was presented in the RVF **(Figure 6)**. Thus, spectral analysis was conducted on the deviation differences between valid and invalid trials. The results showed significant 19 Hz (cue at RVF; permutation test, corrected p < 0.05) and 15 Hz (cue at LVF; permutation test, corrected p < 0.05) oscillations in the 50% cue validity condition **(Figure 7).**

**Figure 4.**
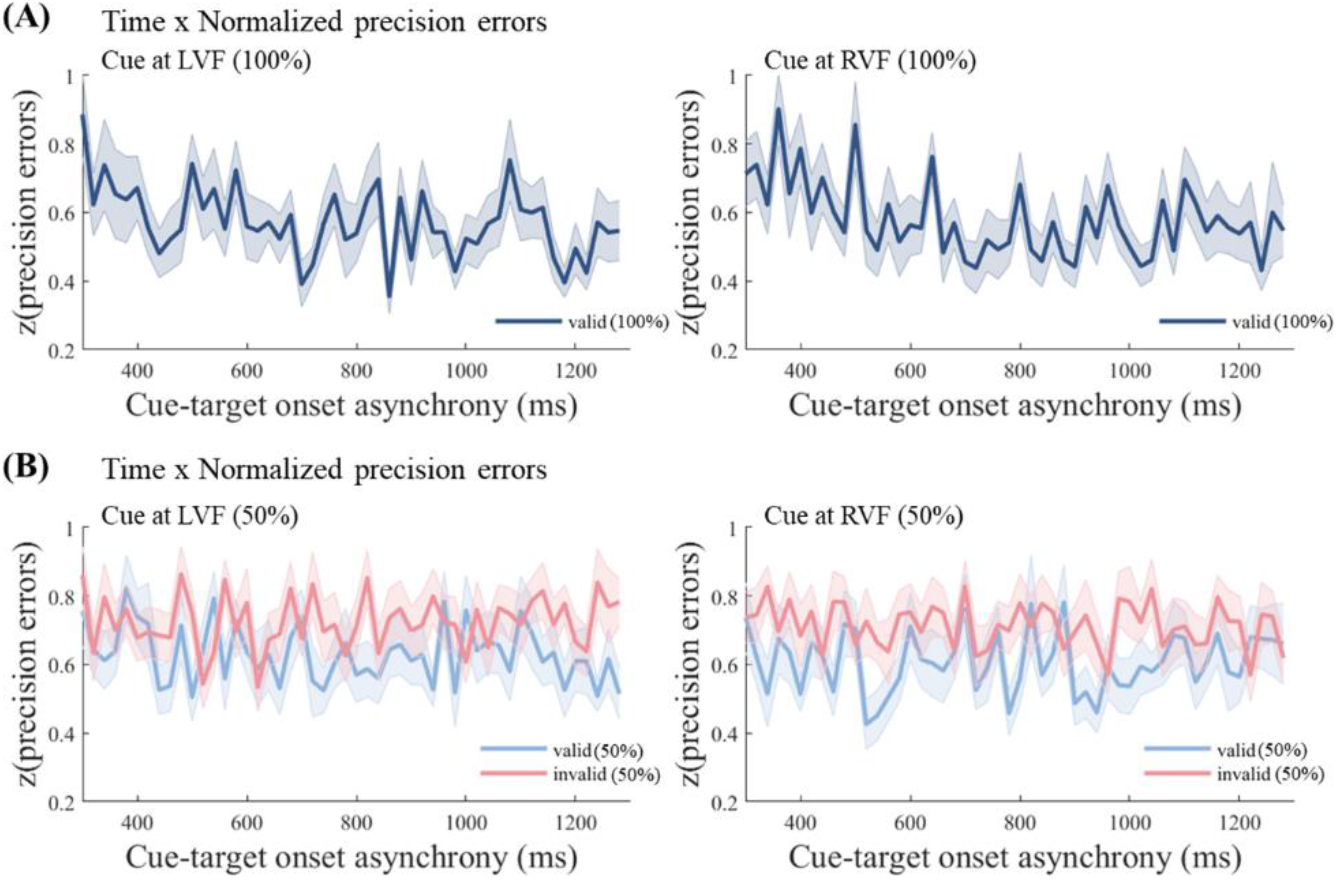
Normalized precision errors alternate as function of CTOA. (A) shows the normalized precision errors in valid trials under the 100% cue validity condition. (B) displays the normalized precision errors in both valid (blue) and invalid (pink) trials under the 50% cue validity condition. Note that in all figures, the left side indicates trials cued at the left visual field (LVF), while the right side indicates trials cued at the right visual field (RVF).

**Figure 5.**
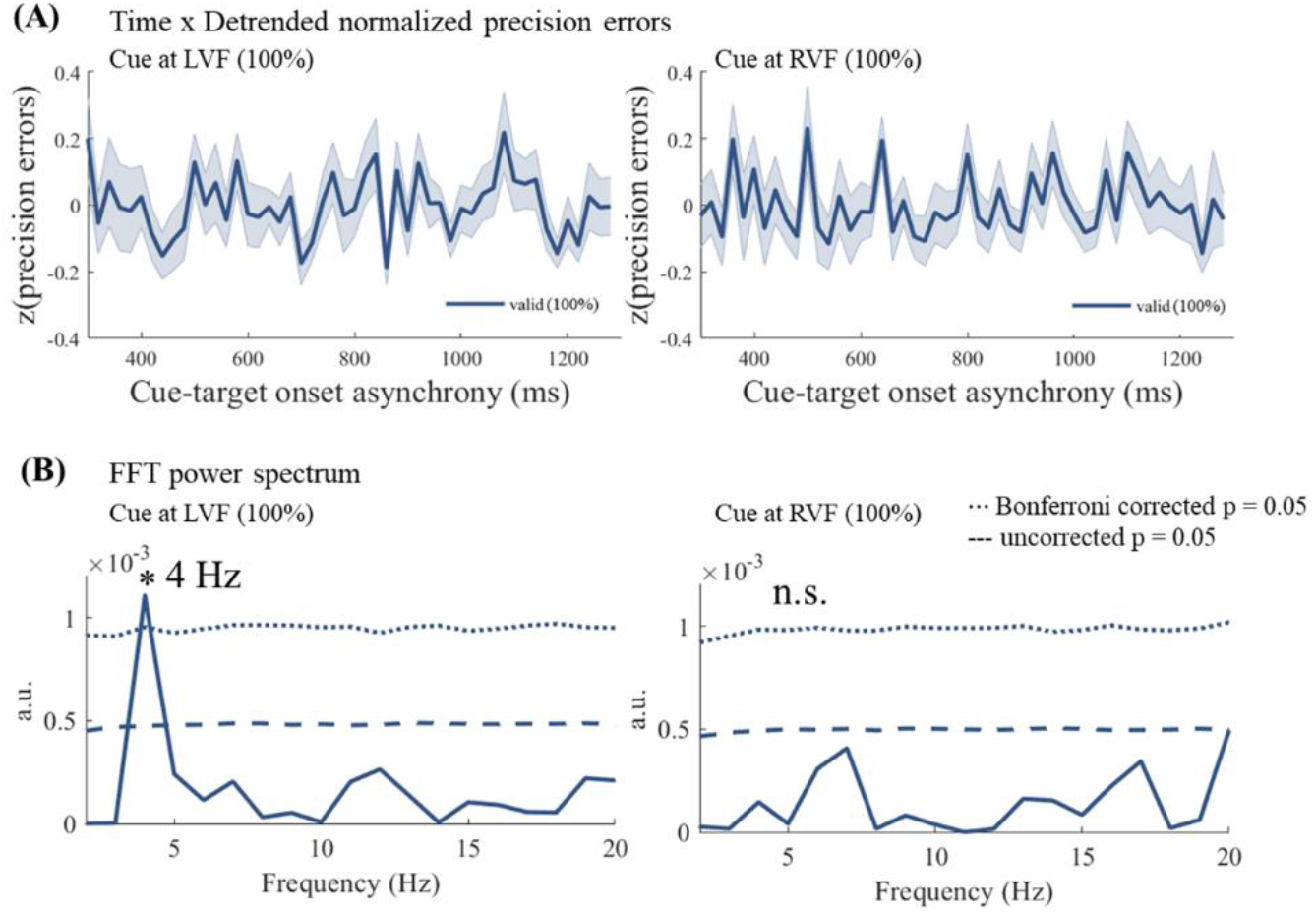
Behavioral fluctuations at theta-band in the 100% cue validity condition. (A) Time-detrended normalized precision errors in 100% cue validity condition. (B) Results of the spectral analysis indicate a significant 4 Hz rhythm in the left visual field (LVF). Solid lines represent power estimates, dashed lines represent a p-value of 0.05 from the permutation test, and dotted lines represent Bonferroni corrected p-value of 0.05. In all figures, the left side represents trials cued at the left visual field (LVF), while the right side represents trials cued at the right visual field (RVF). (* = statistically significant; p< 0.05, n.s. = not statistically significant)

**Figure 6.**
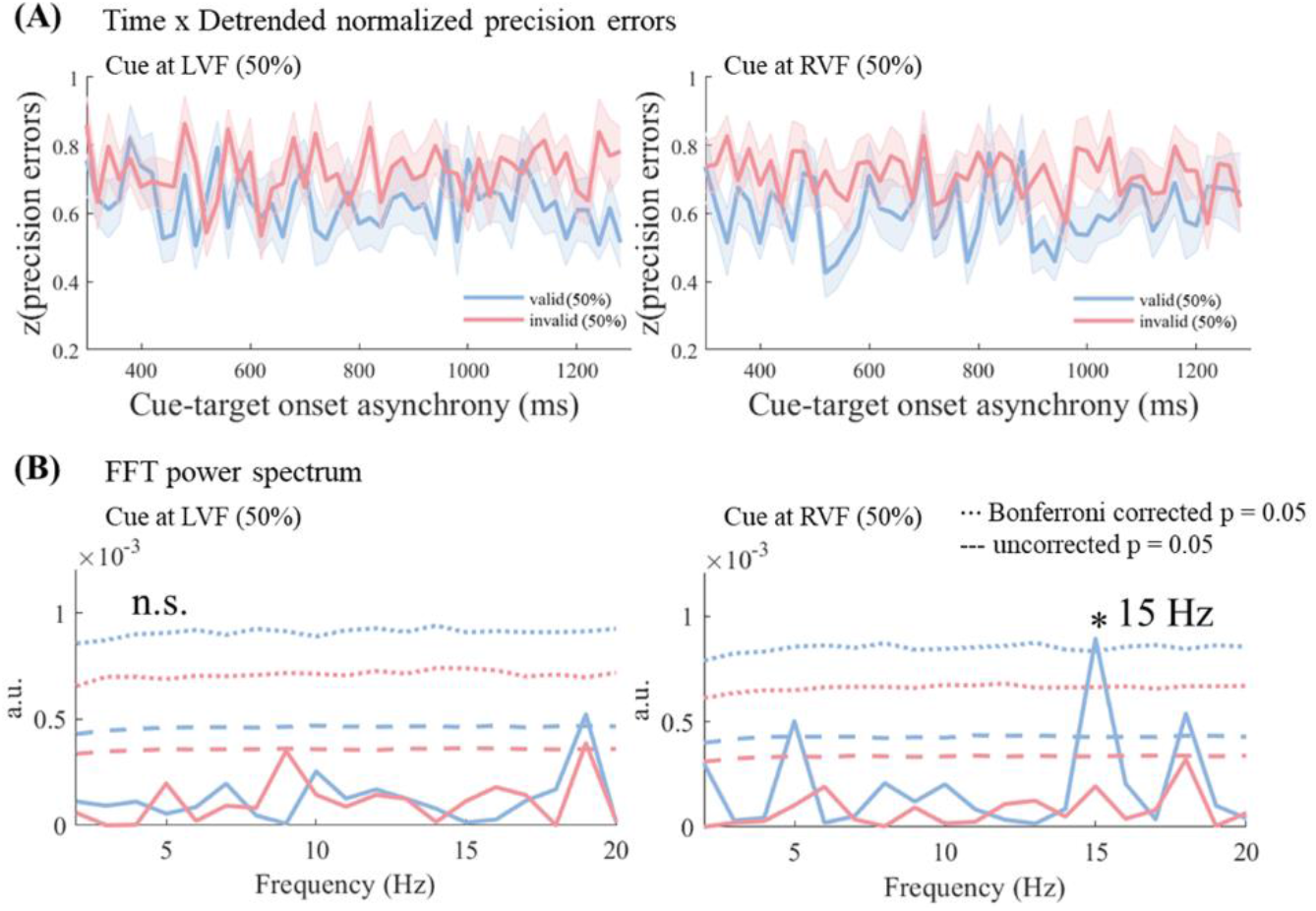
Behavioral fluctuations in 50% cue validity condition. (A) Time-detrended normalized precision errors in the valid (blue) and invalid (pink) trials of 50% cue validity condition are illustrated. (B) The results of the spectral analysis indicate a significant 15 Hz rhythm in valid trials, cued at the right visual field (RVF). Solid lines represent power estimates, the dashed line represents a p-value of 0.05 from the permutation test, and the dotted line represents the Bonferroni corrected p-value of 0.05. Note that in all figures, the left side represents trials cued at the left visual field (LVF), while the right side represents trials cued at the right visual field (RVF). (* = statistically significant (p< 0.05), n.s. = not statistically significant)

**Figure 7.**
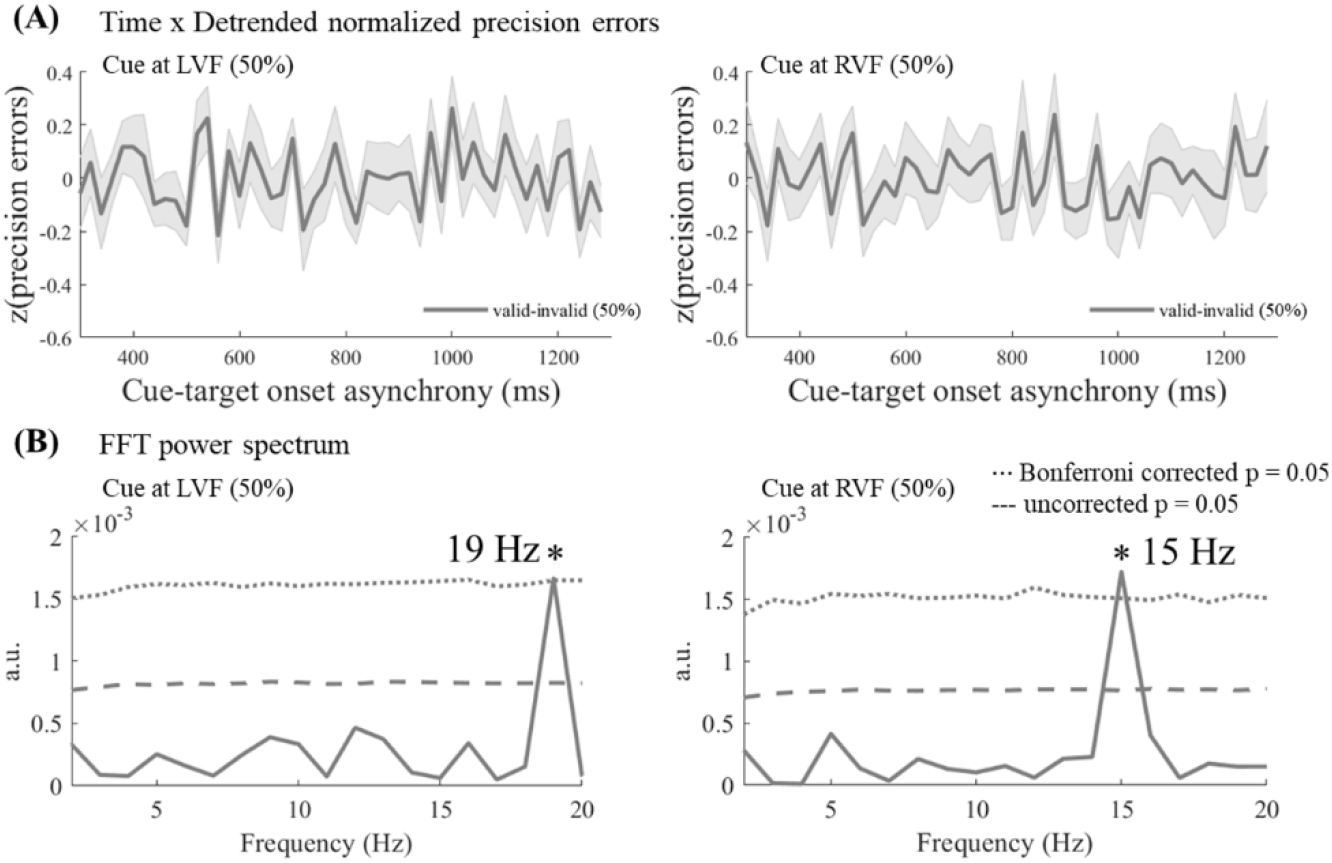
Validity effects on precision errors in 50% cue validity condition. (A) Differences in normalized precision errors between the valid and invalid trials of the 50% cue validity condition are illustrated. (B) The spectral analysis results indicate a significant 19 Hz rhythm cued at the left visual field (LVF) and a significant 15 Hz cued at the right visual field (RVF). Solid lines represent power estimates, the dashed line represents a p-value of 0.05 from the permutation test, and the dotted line represents the Bonferroni corrected p-value of 0.05. Note that in all figures, the left side represents trials cued at the left visual field (LVF), while the right side represents trials cued at the right visual field (RVF). (* = statistically significant (p< 0.05), n.s. = not statistically significant)

### 3.2 Neural Correlates of Spatial Cues

#### 3.2.1 Asymmetric Alpha Oscillations

The neural response to the spatial cue was initially examined using 2D HHS (T-CF & CF-AM frequency). Previous research has shown that spatial attentional modulation leads to lateralized power changes in neural oscillations in parietal-occipital areas, specifically in the alpha-band (Peylo et al. 2021). Our findings from the time-carrier frequency spectrum depicted significant asymmetric alpha oscillations (8-16 Hz) and theta oscillations in the parietal-occipital areas, particularly in the left cluster EEG channels (paired t-test, CBnPP test, cluster p < 0.02, max t = 5.38, 200-300 ms in the alpha-band), indicating the impact of cue validity in the 100% cue validity condition **(Figure 8A)**. However, there were no significant alpha asymmetric findings (paired t-test, CBnPP test, cluster p = 0.09) in the 50% cue validity condition **(Figure 8B)**. Furthermore, in line with recent studies suggesting the involvement of phase coupling between low-frequency waves and amplitude modulation of high-frequency waves within the visual system during attentional modulation (Chacko et al. 2018), further analysis was conducted on the frequency of amplitude envelopes derived from lateralized neural oscillations using the multiple layer HHSA method (Nguyen et al. 2019). As a result, our analysis of amplitude modulation during the preparatory period revealed that the asymmetric alpha oscillations (8- 16 Hz) were modulated by the low-frequency band (1-4 Hz) in left occipital clusters (paired t-test, CBnPP test, cluster p = 0.006, within the theta-modulated alpha range) in the 100% cue validity condition **(Figure 9A)**. However, only marginal significance was observed in right occipital clusters (paired t-test, CBnPP test, cluster minimum p = 0.05) in the 50% cue validity condition **(Figure 9B)**.

**Figure 8.**
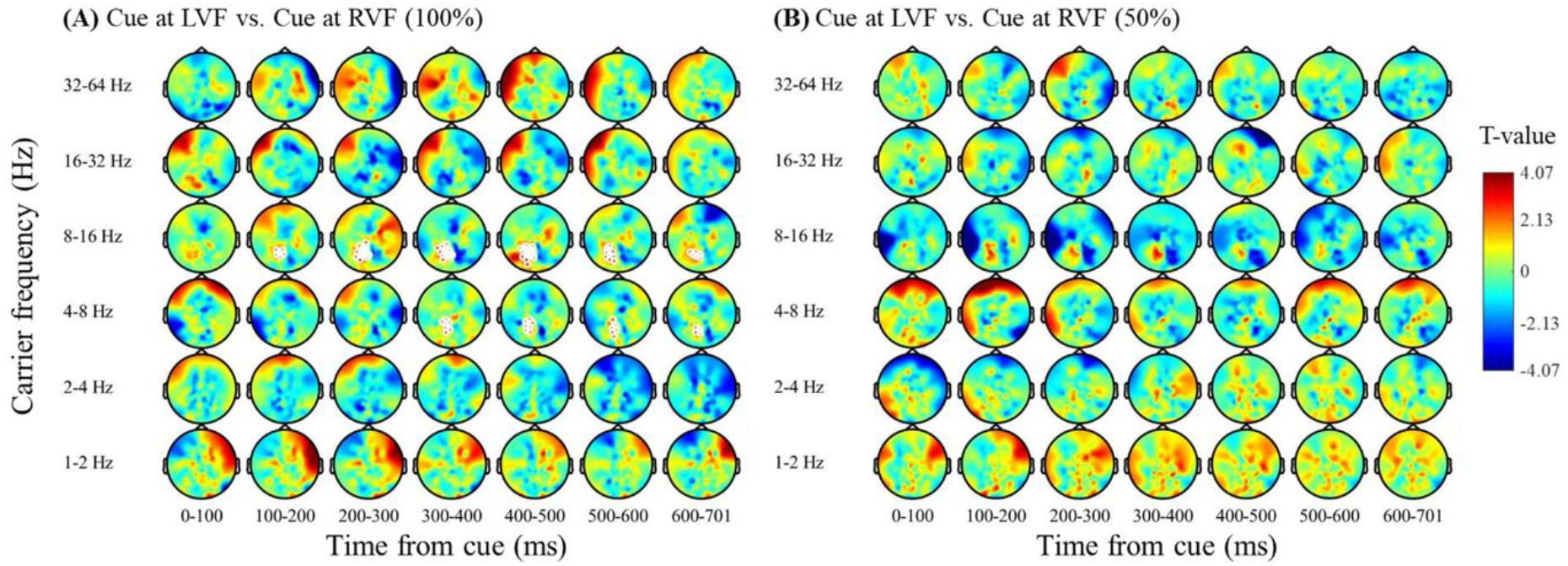
The time-Carrier frequency spectrum of spatial attention modulation. The figure illustrates the results of pair-t tests comparing the rescaled power between trials cued at the left visual field (LVF) and right visual field (RVF) in both the 100% cue validity condition (shown in A) and the 50% cue validity condition (shown in B) within the time window of 0-700 ms after the exogenous cue. The white markers indicate significant differences detected by the paired t-CBnPP test (two-tailed, p < 0.05). Notably, significant alpha lateralization was observed in the left parietal-occipital clusters, specifically in the 100% cue validity condition.

**Figure 9.**
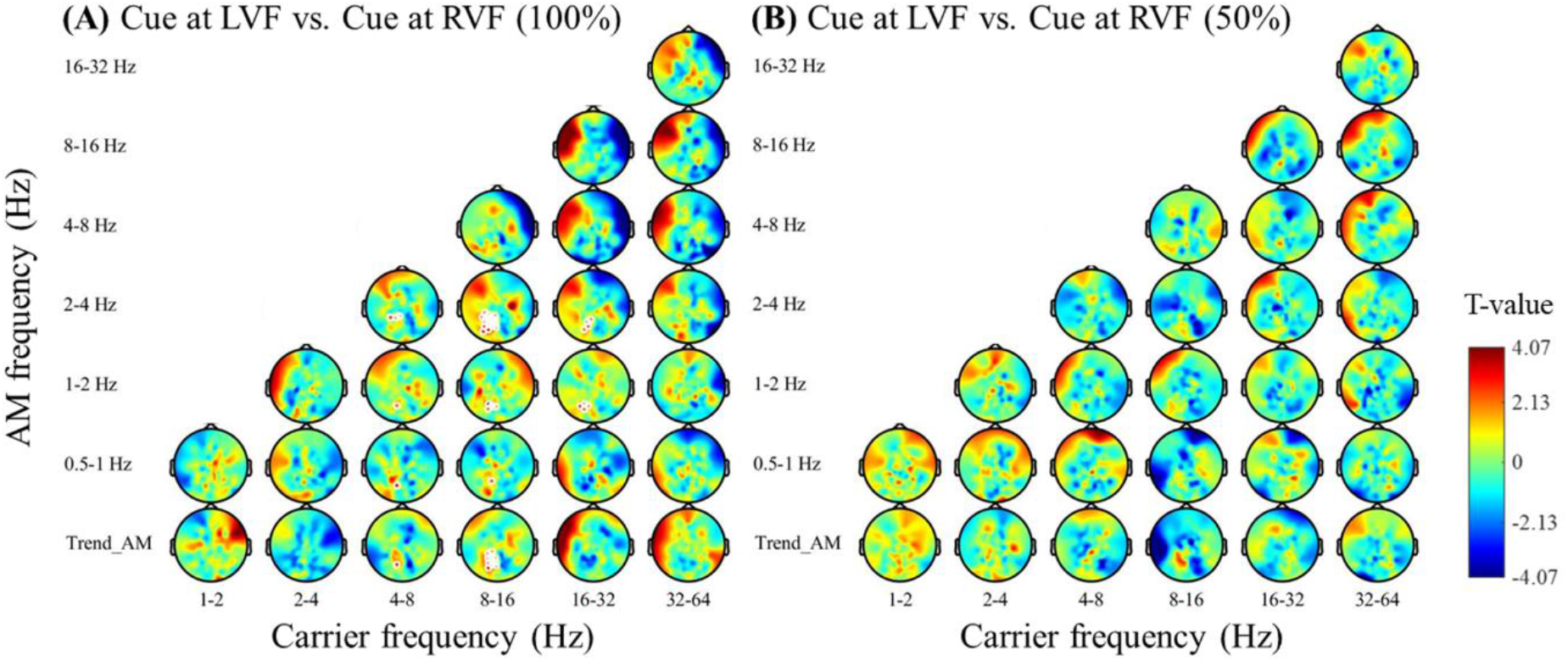
Carrier frequency-Amplitude modulation spectrum of spatial attention modulation (0 to 700ms from the cue onset). The figure presents the results of pair-t tests comparing the rescaled power between trials cued at the left visual field (LVF) and right visual field (RVF) in both the 100% cue validity condition (depicted in A) and the 50% cue validity condition (depicted in B) within the time window of 0-700 ms after the exogenous cue. The white markers indicate significant differences detected by the paired t-CBnPP test (two-tailed, p < 0.05). Notably, in the 100% cue validity condition, a modulation of alpha amplitude by low frequencies (1-4 Hz) was observed.

### 3.3 Cue Validity Effects on Neural Communications Inter-areal Phase Amplitude Coupling

Based on the observation of low-frequency amplitude modulations of the alpha activities at the parietal-occipital areas, the Holo-Hilbert cross-frequency phase clustering (HHCFPC) was conducted aiming to search for the origin of low-frequency modulations and their interactive features. The seed of the alpha-range receiver (IMF5 & 6: 5.7-19.9 Hz) was first set at right and left occipital clusters. As a result, as illustrated in **Figure 10A**, it was found that low frequency (IMF7: 2.6-5.7 Hz) phase of left frontal areas has stronger connectivity with high frequency (IMF5& 6: 5.7-19.9 Hz) amplitude function at the right parietal-occipital clusters (paired t-test, CBnPP test, sender cluster p = 0.046, maximum t = 4.75 at E28) in 100% cue validity condition. However, there was no difference between cue validities while placing the receiver at the left parietal clusters. To further examine the role of low frequency at the frontal areas, we further place the sender (IMF7: 2.6-5.7 Hz) seed to the most significant channel at the left frontal area (E28) and the right frontal area (E117, symmetric to E28). The results showed that the theta phase/ alpha-beta amplitude coupling was significantly larger in the 100% cue validity condition **(Figure 10B)**. Both the theta oscillations at the right and left frontal areas were phase-coupled with the alpha-beta activities at right parietal-occipital areas from 100 ms after cue onset (paired t-test, CBnPP test, right frontal seed: p = 0.004, left frontal seed: p = 0.016). Furthermore, the HHCFPC was found to enhance within the frontal areas in 100% cue validity condition (paired t-test, CBnPP test, right frontal seed: p = 0.004, left frontal seed: p = 0.017). On the other hand, we also examined the difference in theta phase/beta-gamma amplitude coupling between different cue validity conditions. Similar to the previous analysis, the sender (IMF7: 2.6-5.7 Hz) was placed at the right (E117) and left (E28) frontal channels, while the receiver was selected from the beta-gamma frequency range (IMF4: 19.9-43.3 Hz). However, no significant difference was found between the cue validity conditions, regardless of the right (paired t-test, CBnPP test, receiver cluster p = 0.17) or left (paired t-test, CBnPP test, receiver cluster p = 0.13) frontal sender (**Figure 11**).

**Figure 10.**
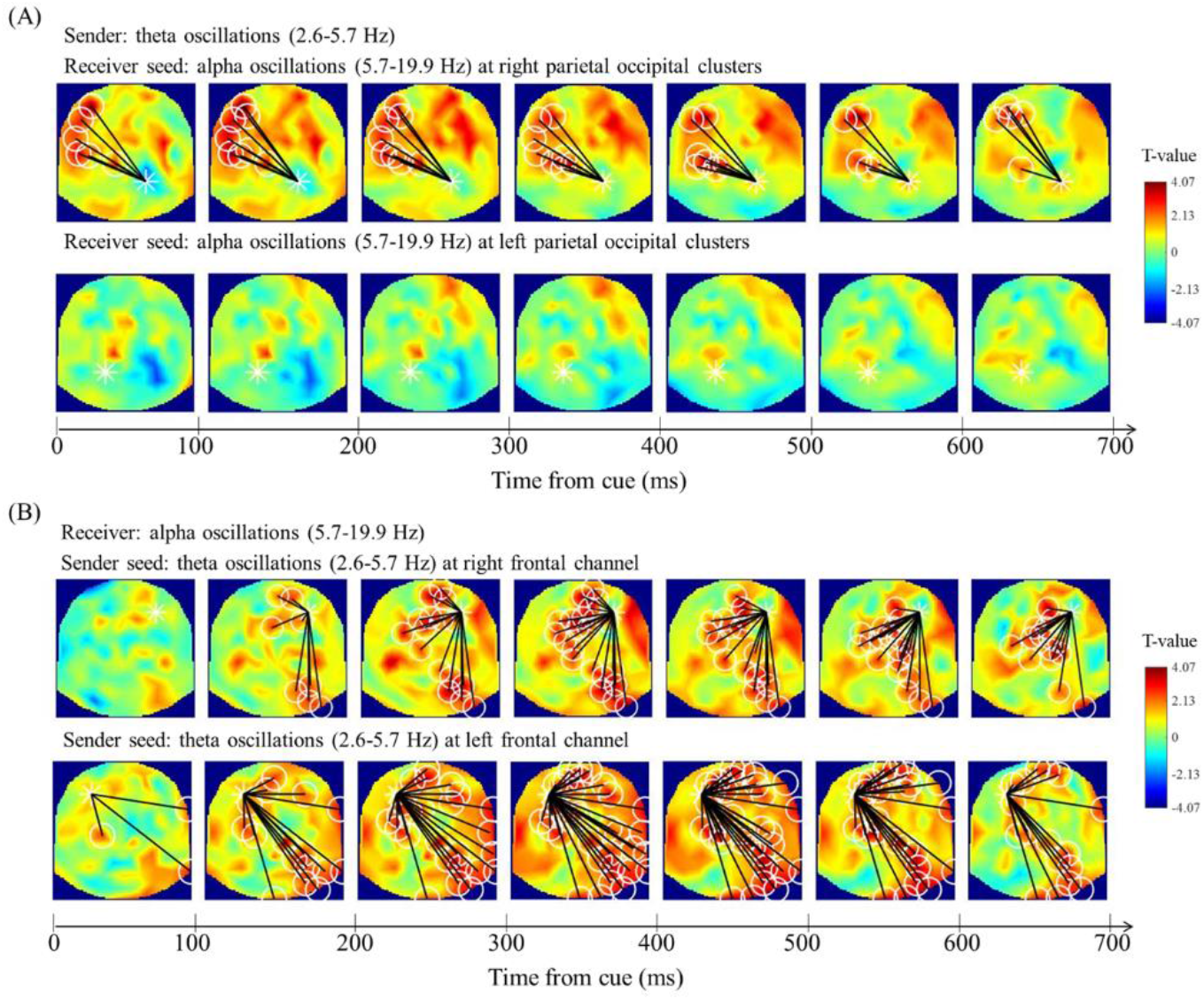
Effects of cue validity on theta phase/alpha amplitude HHCFPC. The figure illustrates the difference in theta phase/alpha amplitude Holo-Hilbert Cross-frequency Phase Clustering (HHCFPC) between cue validity conditions from 0-700 ms after cue onset. The receiver seed (IMF5 & 6: 5.7-19.9 Hz) was placed in the right (A upper) and left (A lower) parietal-occipital clusters. Additionally, the sender seed (IMF7: 2.6-5.7 Hz) was placed in the right (B upper) and left (B lower) frontal channels. The white markers indicate significant differences detected by the paired t-CBnPP test (two-tailed, p < 0.05). Notably, stronger interareal (frontal and right parietal) and within frontal theta phase/alpha amplitude HHCFPC was found in the 100% cue validity condition.

**Figure 11.**
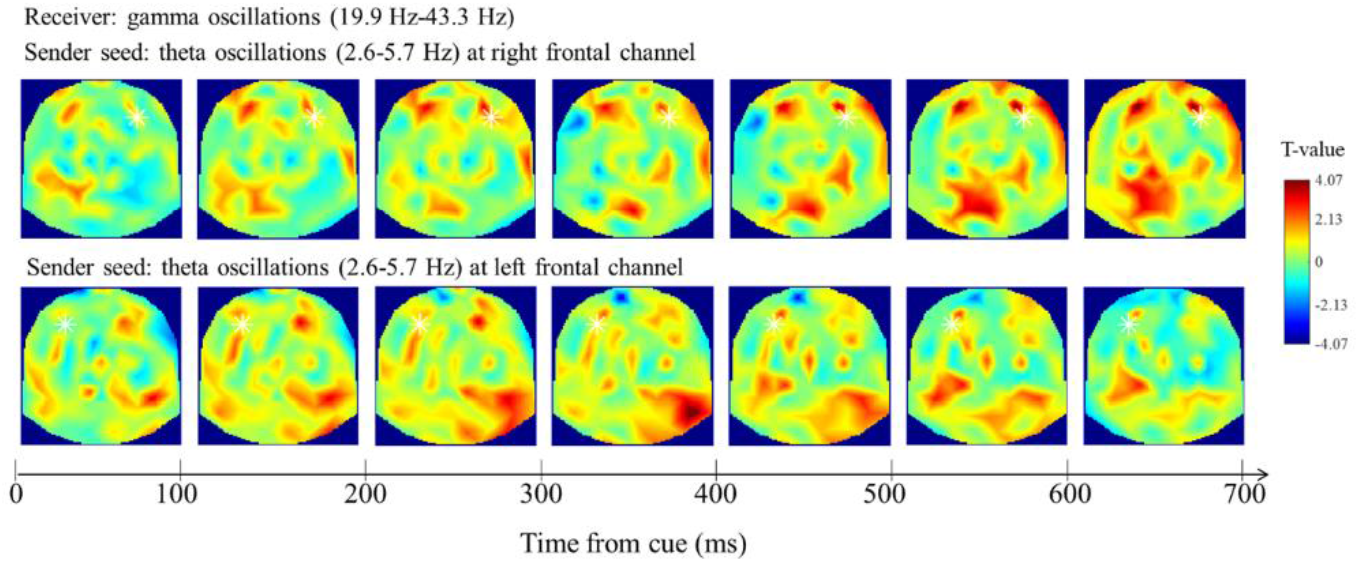
Effects of cue validity on theta phase/gamma amplitude HHCFPC. The figure illustrates the difference in theta phase/gamma amplitude Holo-Hilbert Cross-frequency Phase Clustering (HHCFPC) between cue validity conditions from 0-700 ms after cue onset. The sender seed (IMF7: 2.6-5.7 Hz) was placed in the right (upper) and left (lower) frontal channels, while the receiver was selected from the beta-gamma frequency range (IMF4: 19.9-43.3 Hz). There was no significant difference in theta phase/gamma amplitude HHCFPC between cue validity conditions.

### 3.4 Neural Correlates Preceding the Target

#### 3.4.1 Spectral Power Predicts Response Precision

In previous studies, the power of neural oscillations in the pre-stimulus period was thought to be predictable for the subsequent perception-related performance, particularly at the alpha-band (Mathewson et al. 2009; Michail et al. 2022). This study first examined the pre-stimulus period through the T-CF frequency spectrum. The paired t-test, CBnPP test was conducted between the good performance phase (i.e., precise) and the bad performance phase (i.e., imprecise) within each condition. As a result, there was no significant power difference before the stimuli onset in the 100% cue validity condition (paired t-test, CBnPP test, minimum p = 0.19, **Figure 12A**). However, gamma oscillations, instead of alpha oscillations, at parietal-occipital areas were significantly higher in precise response compared to the imprecise response in the 50% cue validity condition during the pre-stimuli period (-700-0 ms, paired t-test, CBnPP test, p = 0.02, **Figure 12B**). To further examine the amplitude functions of gamma oscillations at parietal-occipital areas, the CF-AM spectrum was calculated during the pre-stimuli period (-700-0 ms). As a result, in occipital clusters (O1-OZ-O2: E70, E71, E74, E75, E76, E82, E83) beta (16-30 Hz) modulating high gamma activities (paired t-test, CBnPP test, p = 0.024) and alpha (7-8 Hz) modulating low gamma activities (paired t-test, CBnPP test, p = 0.042) were found significant between precise and imprecise response in the 50% cue validity condition (**Figure 13A**), with no differences in the 100% cue validity condition in the pre-stimuli period (**Figure 13B)**.

**Figure 12.**
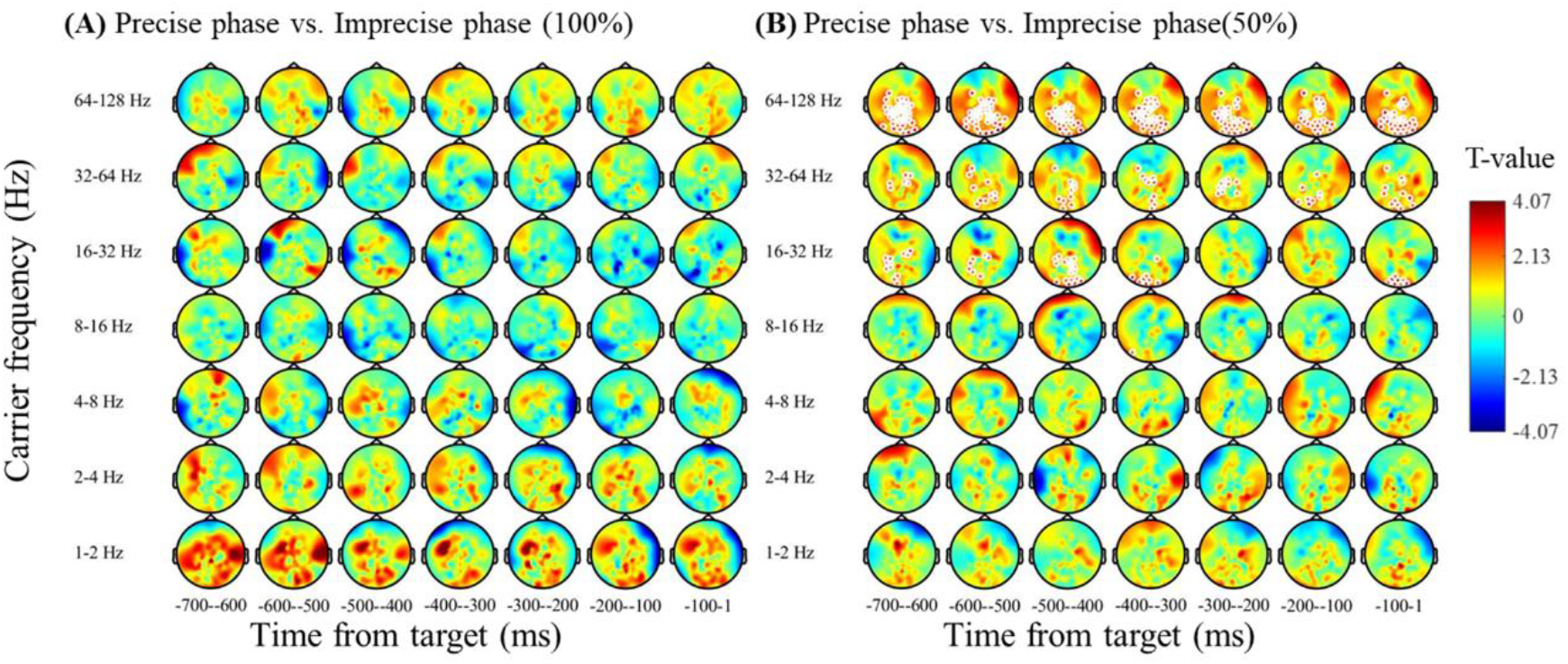
Pre-stimulus neural oscillations predict response precision (T-CF). The figure illustrates the results of paired-t tests comparing the rescaled power between precise and imprecise trials in both the 100% cue validity condition (shown in A) and the 50% cue validity condition (shown in B) within the time window of -700-0 ms before the target. The white markers indicate significant differences detected by the paired t-CBnPP test (two-tailed, p < 0.05). Notably, significant beta-gamma oscillations (> 16 Hz) were observed in the parietal-occipital clusters, specifically in the 50% cue validity condition.

**Figure 13.**
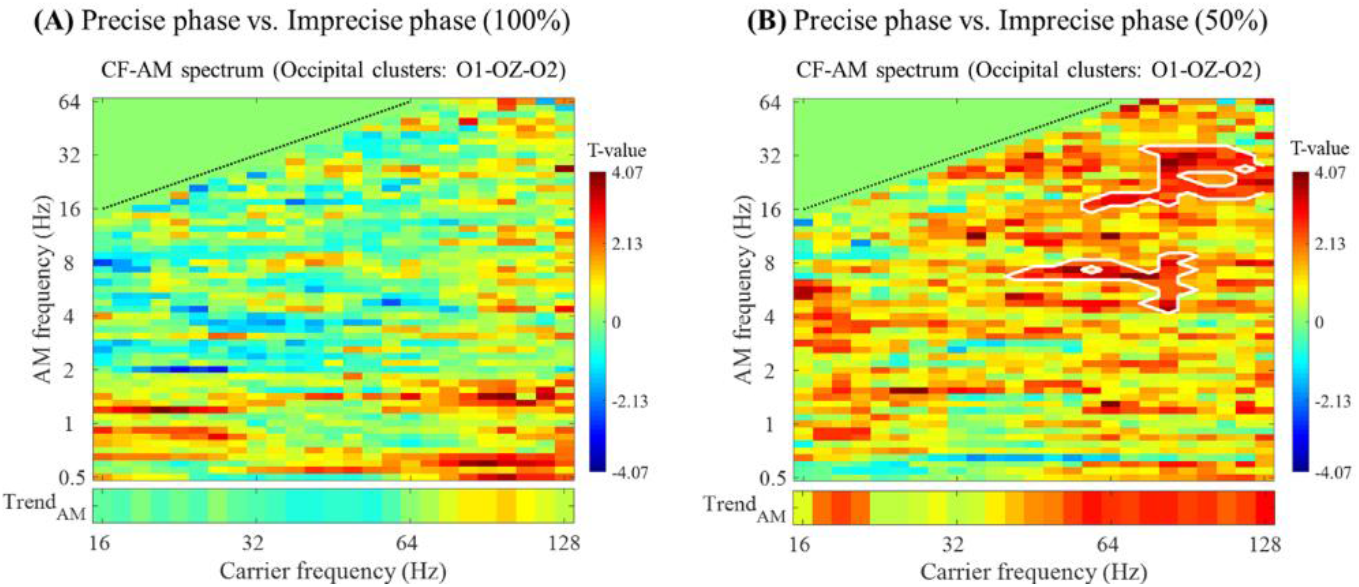
Occipital beta-modulated gamma activities predict response precision (CF-AM). The figure shows the carrier frequency-amplitude modulation spectrum depicting the statistical difference between precise and imprecise trials within the 100% (A) and 50% (B) cue validity conditions at the occipital clusters, averaged from -700 ms before target onset to target onset. The white boxes highlight the areas that show statistical significance based on the two-tailed CBnPP test (p<0.05). Notably, alpha-beta modulated gamma activities were found to predict the response precision phase in the 50% cue validity condition.

#### 3.4.2 Inter-areal Phase Amplitude Coupling Predicts Response Precision

Following the analysis within the preparatory period, we also examined whether theta-phase/alpha-beta amplitude coupling or beta-phase/gamma amplitude coupling is predictable for behavioral performance. Subsequent to the previous analysis, the sender (IMF7: 2.6-5.7 Hz) seed was placed at the left (E28) and right (E117) frontal channels. In the comparison of precision and imprecise trials of 100% cue validity conditions **(Figure 14A**), we found a significant difference (paired t-test, CBnPP test, p = 0.016, max t = 5.10) between the HHCFPC of theta phase at right frontal areas and alpha-beta amplitude at occipital-parietal areas 400- 600 ms before the stimuli onset. However, there was no difference between the precise and imprecise trials in the 50% cue validity condition (**Figure 14B**).

**Figure 14.**
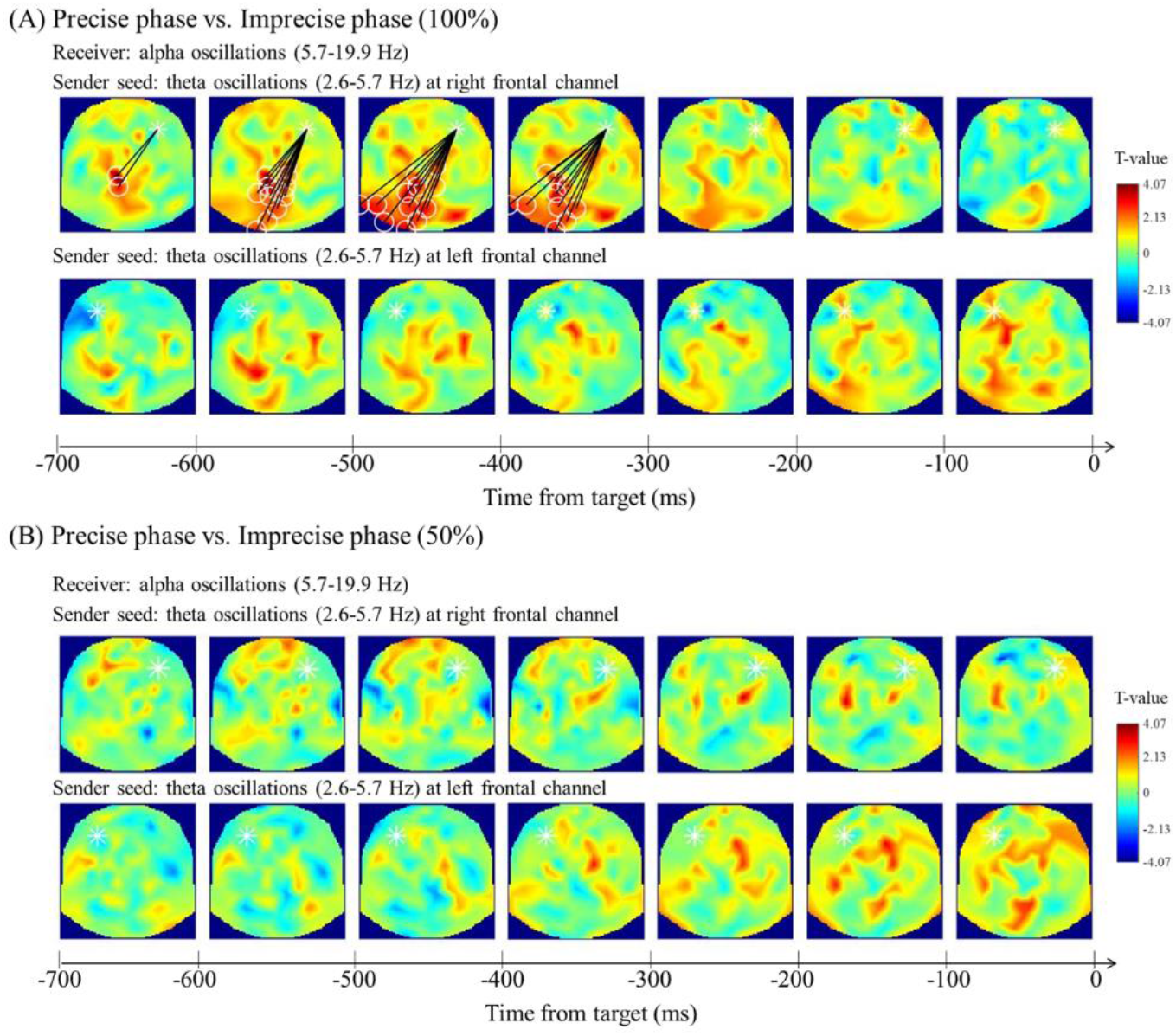
Frontal theta phase/occipital alpha amplitude HHCFPC predicts response precision. The figure illustrates the difference in theta phase/alpha amplitude Holo-Hilbert Cross-frequency Phase Clustering (HHCFPC) between precise and imprecise trials within 100% and 50% cue validity conditions from 0-700 ms after cue onset. The receiver seed (IMF5 & 6: 5.7-19.9 Hz) was placed in the right (A upper) and left (A lower) parietal-occipital clusters. Additionally, the sender seed (IMF7: 2.6-5.7 Hz) was placed in the right (B upper) and left (B lower) frontal channels. The white markers indicate significant differences detected by the paired t-CBnPP test (two-tailed, p < 0.05). Notably, stronger interareal (frontal and right parietal) and within frontal theta phase/alpha amplitude HHCFPC was found in the 100% cue validity condition.

## 4. Discussion

This study aimed to explore the temporal dynamics of sustained spatial attention under varying levels of certainty, utilizing behavioral and electrophysiological measures in an adaptive discrimination task. The behavioral analysis results uncovered distinct attentional effects induced by certain and uncertain spatial cues on perceptual sensitivity. Specifically, our findings indicated that an increase in spatial certainty leads to a decrease in attentional sampling rhythms from the beta-band to the theta-band. Furthermore, the EEG results provided insights into the underlying neural mechanisms, showcasing diverse activations and functional communications, particularly through theta phase/alpha amplitude coupling, within the attentional system involving the frontal-parietal network and the visual cortex. In summary **(Figure 15)**, the study identified synergistic effects between spatial and temporal prediction and contributed to existing knowledge by demonstrating that the certainty of spatial prediction modulates attentional sampling rhythms.

**Figure 15.**
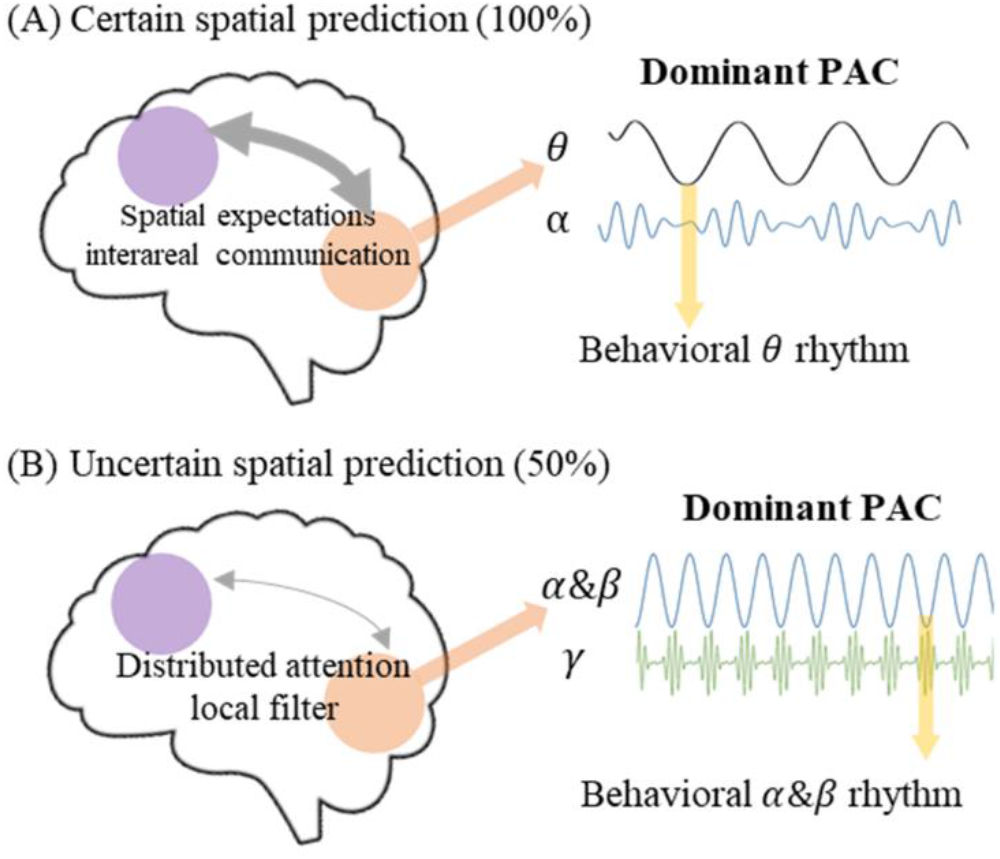
Certainty of spatial cue impact on brain and behavioral rhythms (A) In the left figure, certain spatial cues trigger strong spatial expectations by enhancing interareal communication between the frontal (purple) and parietal-occipital (orange) areas. The right figure illustrates the dominant theta phase/alpha amplitude coupling within the frontal-parietal network during specific spatial predictions, which prevails in influencing the behavioral rhythm at the theta frequency. (B) The left figure demonstrates that uncertain spatial cues lead to distributed attention and result in weak interareal modulations. The right figure shows the dominance of the local filtering mechanism of alpha & beta phase/gamma amplitude modulation within the early visual cortex under uncertain spatial predictions. This mechanism, which remains unaffected by top-down modulation, plays a significant role in behavioral rhythms at alpha and beta frequencies.

Our findings suggest that spatial cues indeed influence perceptual sensitivity, and different levels of certainty in spatial cues contribute to varying levels of preparation for target perception. We observed that certain spatial predictions led to improved discrimination of the small gap in the near-threshold target (i.e., Landolt ring) compared to uncertain spatial predictions, aligning with previous studies indicating that prior expectations can enhance perceptual sensitivity (e.g., Bashinski and Bacharach 1980). Interestingly, we also noted cueing effects even under uncertain spatial prediction, as evidenced by significant differences in precision errors between valid and invalid trials in the 50% cue validity condition.

Moreover, our study uncovered evidence of distinct fluctuations in precision errors, indicating that spatial certainty modulates attentional sampling rhythms. Specifically, we observed theta (4 Hz) oscillations of precision errors in the 100% cue validity condition. In contrast, alpha & beta (15 & 19 Hz) oscillations of precision errors were present in the 50% cue validity condition. Expanding upon previous research suggesting that inter-areal cross-frequency coupling contributes to theta-phasic effects on behavioral performance (Fiebelkorn et al. 2018; Plöchl et al. 2022), we analyzed phase-amplitude coupling within the activated attentional network and visual cortex. As mentioned earlier, our findings revealed robust frontal-parietal and within- frontal theta phase/alpha amplitude coupling induced by certain spatial cues, with frontal- parietal theta phase/alpha amplitude coupling in the pre-target period predicting response precision in the 100% cue validity condition. Additionally, intra-areal beta modulation of gamma activities within the occipital areas predicted response precision in the 50% cue validity condition. In summary, we found that behavioral fluctuations were reflected in the phase of amplitude modulation within the visual system, aligning with the current model suggesting that sampling rhythms are related to task-induced neural oscillations (Dugué and VanRullen 2017; Kawashima et al. 2022; Michel et al. 2022).

While our findings support the model proposing that attentional sampling rhythms are linked to functional communications within the visual system, the underlying functions of observed rhythms differ slightly from the current understanding of attentional sampling rhythms and our expectations. Initially, we expected that sequential sampling between two locations would occur around theta frequency, revealing efficient rhythmic search (Fiebelkorn and Kastner 2019), while periodic sampling within one location would occur around alpha frequency, indicating a natural filter (Dugué and VanRullen 2017; Michel et al. 2022) . However, in our study, when we manipulated cue validity to guide participants to sample specifically within one or two locations, we obtained results contrary to these expectations, highlighting the complexity of attentional sampling behavior. This discrepancy raises questions regarding the functional distinctions between the observed theta rhythms and the attentional rhythms reported in previous studies. In fact, our findings suggest that theta rhythms reflect the modulated effects of spatial-temporal expectations in a specific spatial cue condition, indicating long-range communication from higher-order areas, namely the prefrontal cortex, to the visual cortex (Cavanagh and Frank 2014; Paneri and Gregoriou 2017). On the other hand, beta rhythms represent the local sampling mechanism within the early visual cortex in an uncertain spatial cue condition, reflecting local processing and the requirement of distributed attention in the absence of certain spatial cues (Jensen and Mazaheri 2010).

Additionally, our results revealed a hemispheric difference in both behavioral rhythms and inter-areal phase-amplitude coupling, specifically in the condition with certain spatial prediction. We observed behavioral rhythms of response precision at 4 Hz when the cue was presented at the LVF under certain spatial prediction, whereas no specific fluctuations were observed at the RVF. Moreover, our results consistently indicate activation of the right parietal- occipital cortex during the task, regardless of the cued visual fields. This finding is further supported by the observed inter-areal phase-amplitude coupling, where both the right and left frontal eye fields (FEF) exhibited stronger coupling with the right parietal-occipital lobes (Bressler et al. 2008; Grosbras and Paus 2003; Siman-Tov et al. 2007). Additionally, lateralized alpha modulation was only observed in the left parietal-occipital lobes. Previous studies have highlighted the crucial role of the right parietal-occipital cortex in spatial attention, including the maintenance of spatial information (Malhotra et al. 2009), reorienting of spatial attention (Chang et al. 2016; Chang et al. 2013) discrimination of spatial orientation in clock angles (Sack 2009), and rhythmicity of sampling mechanism between two visual fields (Landau and Fries 2012). Therefore, in the condition where the cue is presented in the RVF, it was suspected that the simultaneous presence of non-target stimuli (i.e., distractors) in the LVF could influence target encoding in the RVF, as the corresponding right parietal-occipital lobes were activated. To uncover the underlying sampling mechanisms responsible for these hemispheric differences, future studies using non-invasive brain stimulation methods like transcranial alternating current stimulation (tACS) with rhythmic entrainment may provide a causal investigation into the specific roles of these brain regions and the impacts of modulation (e.g., Kasten et al. 2020; Tseng et al. 2016).

Although our results showed a significant correlation between response precision and pre- target neural correlates, the experimental design of delayed response raises the possibility that preparation is not limited to attentional processes but also involves working memory. The process of working memory involves several phases, including encoding, maintenance, and retrieval. Recent research has shed light on the role of low-frequency oscillations (2-13 Hz) within the cortex in memory encoding and retrieval (Abdalaziz et al. 2023; Cruzat et al. 2021; Mohan et al. 2022). Specifically, a previous study suggested that theta and beta phases of brain oscillations in the frontal and parietal cortex serve as mechanisms to prevent representation conflict during the memorization of multiple items (Abdalaziz et al. 2023). This suggests that the preparation of working memory involves the allocations of neural oscillations to optimize encoding and retrieval processes. It is intriguing to note that the sampling rhythms observed in our study, particularly under conditions of uncertain spatial prediction, align with the beta band, which is associated more commonly with working memory, rather than the typical association of the alpha band with perceptual rhythms (VanRullen 2016). This piece of evidence suggests that it is possible that attention and preparation of encoding share the same resources, such as the frontal-parietal network and visual system (Balestrieri et al. 2022). These shared resources likely contribute to the modulation of sampling rhythms, influencing the efficiency of cognitive functions.

Recent studies have introduced various experimental paradigms to investigate how specific task demands interact with the rhythm of periodic sampling. As aforementioned, studies applied dual tasks found that the phasic contributions of other cognitive functions can influence target perception, such as working memory (Balestrieri et al. 2022) and motor actions (Tomassini et al. 2017). Moreover, it is suspected that task parameters such as the number of sampling objects (Fiebelkorn et al. 2013; Holcombe and Chen 2013; Re et al. 2019), occurrence of distractors (Kawashima et al. 2022), task difficulties (Chen et al. 2017; Merholz et al. 2022), rewards (Su et al. 2021), and continuous response (Michel et al. 2022; Song et al. 2014) can influence the observed rhythms. In summary, our findings contribute to bridging the gap by demonstrating that the validity of spatial cues also influences sampling behavior.

On the other hand, there have been debates in recent studies regarding the existence of phasic contributions of neural oscillations on behavioral performance, despite consistent findings of perceptual and attentional rhythms in several replications. A review of rhythms in cognition highlighted that periodic sampling is not continuously observed in behavioral or electrophysiological studies (Keitel et al. 2022). For instance, the phasic effects of alpha oscillations have sometimes been attenuated from phase-amplitude trade-offs (Fakche et al. 2022; Mathewson et al. 2009), and decreases in alpha-band activities have often been accompanied by decreases in alpha phase/gamma amplitude coupling (Osipova et al. 2008). Some studies have also found evidence for correlations between pre-stimulus alpha power but not phase with target detection (Benwell et al. 2017; Van Dijk et al. 2008). Additionally, null reports from behavioral studies have shown potential instability in the phasic contributions of neural oscillations on behavioral performance, including studies involving endogenous modulations (e.g., van Der Werf et al. 2022) and rhythmic entrainments (Lin et al. 2022).

## 5. Conclusion

In conclusion, our study supported the notion of periodic attentional sampling with converging behavioral and electrophysiological evidence during the preparatory period. These findings shed light on the interplay between certain and uncertain spatial cues and subsequent attentional processes leading up to target onset. Specifically, our study addressed the inconsistent findings in previous research regarding the impact of cue validity on periodic behavioral rhythms. We observed that certain prior spatial expectations decrease the speed of attentional sampling rhythms from the beta to theta band, as supported by consistent behavioral and electrophysiological evidence. Moreover, the certainty of spatial prediction in discriminating ring gaps was associated with distinct functional connectivity patterns in the attentional network. This provides insights into how the brain adapts during different levels of spatial attention certainty in the context of temporally varying targets. These results, in line with current theories, suggest that rhythmic sampling behavior is highly task-dependent. The observed neural rhythms likely serve as a mechanism for the brain to allocate resources adaptively, synchronize neural activities, and enhance the efficiency of information processing and integration during attentional tasks. Overall, our study not only contributes to existing models of attentional sampling rhythms, but expands our understanding of the complex interplay between spatial and temporal predictions in attentional tasks. It paves the way for future research to delve into the intricacies of the brain’s adaptability and explore potential applications in cognitive enhancement, rehabilitation, and brain-computer interface.

## Acknowledgments

We are grateful to Philip Tseng, Wen-Sheng Chang and Yan-Hsun Chen’s comments and suggestions on this work.

